# Anti-microbiota vaccines modulate the tick microbiome in a taxon-specific manner

**DOI:** 10.1101/2021.05.12.443756

**Authors:** Lourdes Mateos-Hernández, Dasiel Obregon, Alejandra Wu-Chuang, Jennifer Maye, Jeremie Bornères, Nicolas Versillé, José de la Fuente, Sandra Díaz-Sánchez, Luis G. Bermúdez-Humarán, Edgar Torres-Maravilla, Agustín Estrada-Peña, Adnan Hodžić, Ladislav Šimo, Alejandro Cabezas-Cruz

## Abstract

Anti-tick microbiota vaccines have been shown to impact tick feeding but its specificity has not been demonstrated. In this study we aimed to investigate the impact of immune targeting of keystone microbiota bacteria on tick performance, and tick microbiota structure and function. Vaccination against *Escherichia coli*, the selected keystone taxon, increased tick engorgement weight and reduced bacterial diversity in *Ixodes ricinus* ticks compared to those that fed on mice immunized against *Leuconostoc mesenteroides*, a non-keystone taxon or mock-immunized group. The abundance of *Escherichia-Shigella*, but not *Leuconostoc* was significantly reduced in ticks fed on *E. coli*-immunized mice and this reduction was correlated with a significant increase in host antibodies (Abs) of the isotype IgM and IgG specific to *E. coli* proteins. This negative correlation was not observed between the abundance of *Leuconostoc* in ticks and anti-*L. mesenteroides* Abs in mice. We also demonstrated by co-occurrence network analysis, that immunization against the keystone bacterium restructure the hierarchy of the microbial community in ticks and that anti-tick microbiota vaccines reduced the resistance of networks to directed removal of taxa. Functional pathways analysis showed that immunization with a live bacterial vaccine can also induce taxon-specific changes in the abundance of pathways. Our results demonstrated that anti-tick microbiota vaccines can modulate the tick microbiome and that the modification is specific to the taxon chosen for host immunization. These results guide interventions for the control of tick infestations and pathogen infection/transmission.

## Introduction

Ticks, like all multicellular eukaryotes, harbor a very diverse group of commensal, symbiotic, and pathogenic microorganisms that collectively comprise the microbiome (1,2). The complex microbial system and ticks share an intimate relationship and this symbiotic association has developed into an essential evolutionary outcome important for tick development, nutritional adaptation, reproductive fitness, ecological plasticity, and immunity (3-6). There is accumulating evidence that non-pathogenic midgut bacteria may also affect tick vector competence and susceptibility to pathogens transmitted by ticks (7-10). The development of high-throughput sequencing technologies and bioinformatics tools in the last decade has significantly improved our knowledge of the phylogenetic and genetic diversity, dynamics, and ecology of the microbial communities in several tick species (2). However, the vast majority of studies of the tick microbiome are restricted to the taxonomic composition, while the functional significance of bacterial community structure and diversity remains largely unexplored (11).

Recent functional metagenomics studies have shown that exploring the taxonomic composition and variability of the tick microbiome underestimates the multidimensional nature of the tick hologenome and that interferences solely based on taxonomic profiles lack biological significance (11-14). Generally, the native tick microbiome is likely composed of bacteria, archaea, fungi, protozoans and viruses with diverse metabolic capacities, which are engaged in a complex network of cooperative and competitive interactions (1,2). Some of these microorganisms, known as keystone species, co-occur with many others and may have a large regulatory effect on the structure, organization, and function of the tick microbiome. The ubiquitousness of the keystone taxa is likely associated with important resources they provide to the overall microbial community and/or the tick host (11,12,14). This suggests that keystones are an essential component of the functional networks and therefore represent ideal targets for the rational manipulation of the microbial composition and function. The functional capacity of the tick microbiome is not equal to the overall number of its individual components, as microbial species strongly and frequently interact with one another and form a complex functional network (14), which can thus be considered as a fundamental unit in microbial communities of ticks. Therefore, the microbial co-occurrence network represents a useful approach to identify the keystoneness of taxa and the potential interactions within the functional networks (15,16). Understanding these microbe-microbe relationships is a critical step for predicting their holistic consequences on tick performance, physiology, and vector competence (13,14,16). In this sense, a recent study by Mateos-Hernández et al., (2020) demonstrated that disturbing the *Ixodes ricinus* microbiome stability by selective immune targeting of ubiquitous and abundant keystone bacteria disrupts the tick-microbiome homeostasis and results in increased mortality of ticks during feeding. This observation concurred with a wide distribution of genes encoding α1,3-galactosyltransferases (α1,3GT) in the *I. ricinus* microbiota and the host’s immune response to α-Gal. Furthermore, immunization with live *Escherichia coli* significantly reduces the relative abundance of Enterobacteriaceae in adult ticks. In this study, we aimed to investigate whether host antibodies (Abs) targeting keystone bacteria could impact the taxonomic and functional profiles of the tick microbiome as well as the structure of the microbial community associated with ticks. Our findings showed for the first time that host immunization with keystone bacteria is a promising tool for experimental manipulation of the tick microbiome composition and activity, and has the potential to reveal other functional mechanisms of the tick-microbiota interactions and could spur new strategies to control ticks and tick-borne pathogens.

## Materials and Methods

### Ethics statement

All procedures were performed at the Animal Facility of the Laboratory for Animal Health of the French Agency for Food, Environmental and Occupational Health & Safety (ANSES), Maisons-Alfort, France, according to French and International Guiding Principles for Biomedical Research Involving Animals (2012). The procedures were reviewed and approved by the Ethics Committee (ComEth, Anses/ENVA/UPEC), with permit number E 94 046 08.

### Mice and housing conditions

Six-week-old C57BL/6 (Charles River strain code 027) mice were purchased from Charles River (Miserey, France) and maintained in Green line ventilated racks (Tecniplast, Hohenpeissenberg, Germany) at −20 Pa, with food (Kliba nafaj, Rinaustrasse, Switzerland) and water *ad libitum*. The mice were kept at controlled room temperature (RT, 20-23°C) and a 12-hour (h) light: 12-h dark photoperiod regimen. The animals were monitored twice a day by experienced technicians and deviations from normal behaviors or signs of health deterioration were recorded, and reported.

### Bacteria cultures and live bacteria immunization

Representative bacteria of the genera *Escherichia*-*Shigella* (i.e., *E. coli*) and *Leuconostoc* (i.e., *L. mesenteroides*) were selected to be included in live bacteria vaccine formulations, aiming to test the impact of host immune response against “keystone” bacteria on tick microbiota composition, stability and functionality, and tick performance. The selection of these bacteria as live vaccines was based on our previous results (16) that show that the family Enterobacteriaceae was among the top keystone taxa (i.e, high relative abundance, ubiquitousness, and eigencentrality) identified in *Ixodes* microbiota. Based on the previous results (16), we selected the family Leuconostocaceae as non-keystone bacteria with low “Keystoneness” (i.e, low relative abundance, ubiquitousness, and eigencentrality) in the microbiome of *Ixodes*.

The gram-negative bacterium *E. coli* BL21 (DE3, Invitrogen, Carlsbad, CA, USA) was prepared as previously described (16). Briefly, *E. coli* was grown on Luria Broth (LB, Sigma-Aldrich, St. Louis, MO, USA) at 37°C under vigorous agitation, washed with phosphate buffer saline (PBS) 10 mM NaH_2_PO_4_, 2.68 mM KCl, 140 mM NaCl, pH 7.2 (Thermo Scientific, Waltham, MA, USA), resuspended at 3.6 × 10^4^ colony-forming unit (CFU)/mL, and homogenized using a glass homogenizer. The gram-positive bacterium *Leuconostoc mesenteroides* (strain LBH1148, INRAE collection) was grown on MRS broth (Difco, Bordeaux, France) at 37°C without agitation and resuspended and homogenized following the same procedures as for *E. coli*. Six-week-old, C57BL/6 mice were immunized subcutaneously with either *E. coli* (n = 4, 1 × 10^6^ CFU per mouse) or *L. mesenteroides* (n = 4, 1 × 10^6^ CFU per mouse) in a water-in-oil emulsion containing 70% Montanide™ ISA 71 VG adjuvant (Seppic, Paris, France), with a booster dose two weeks after the first dose. Control, C57BL/6 (*n* = 4) mice received a mock vaccine containing PBS and adjuvant.

### Bacterial protein extraction

*Escherichia coli* and *L. mesenteroides* were washed twice with PBS, centrifuged at 1000× g for 5 min at 4 °C, resuspended in 1% Trion-PBS lysis buffer (Sigma-Aldrich, St. Louis, MO, USA) and homogenized with 20 strokes using a glass balls homogenizer. The homogenate was then centrifuged at 300× g for 5 min at 4 °C and the supernatant was collected. Protein concentration was determined using the Bradford Protein Assay (Thermo Scientific, San Jose, CA, USA) with Bovine Serum Albumin (BSA) as standard.

### Indirect ELISA

The levels of Abs reactive against bacterial proteins were measured in mice sera as previously reported (16). The 96-well ELISA plates (Thermo Scientific, Waltham, MA, USA) were coated with 0.5 µg/mL (100 µL/well) of *E. coli* or *L. mesenteroides* protein extracts and incubated for 2 h with 100 rpm shaking at RT. Subsequently, plates were incubated overnight at 4 °C. The antigens were diluted in carbonate/bicarbonate buffer (0.05 M, pH 9.6) and incubated overnight at 4 °C. Wells were washed three times with 100 µL of PBS containing 0.05% (vol/vol) Tween 20 (PBST), and then blocked by adding 100 µL of 1% Human Serum Albumin (HSA)/PBS for 1 h at RT and 100 rpm shaking. After three washes, sera samples, diluted 1:50 in 0.5% HSA/PBS, were added to the wells and incubated for 1 h at 37 °C with shaking. The plates were washed three times and HRP-conjugated Abs (goat anti-mice IgG and IgM) (Sigma-Aldrich, St. Louis, MO, USA) were added at 1:1500 dilution in 0.5% HSA/PBST (100 µL/well) and incubated for 1 h at RT with shaking. The plates were washed three times and the reaction was developed with 100 µL ready-to-use TMB solution (Promega, Madison, WI, USA) at RT for 20 min in the dark, and then stopped with 50 µL of 0.5 M H_2_SO_4_. Optimal antigen concentration and dilutions of sera and conjugate were defined using a titration assay. The optical density (OD) was measured at 450 nm using an ELISA plate reader (Filter-Max F5, Molecular Devices, San Jose, CA, USA). All samples were tested in triplicate and the average value of three blanks (no Abs) was subtracted from the reads. The cut-off was determined as two times the mean OD value of the blank controls.

### Immunofluorescence

*Escherichia coli* and *L. mesenteroides* were washed three times with PBS, centrifuged at 1000x g for 5 min, fixed with 4% paraformaldehyde for 30 min and blocked with 1% human serum albumin (HSA, w/v in PBS) for 1h at RT. Bacterial cells were then incubated for two days at 4°C with pooled sera (from all vaccinated mice, day 30) of mice immunized either against *E. coli, L. mesenteroides* or the mock vaccine at a dilution of 1:20 (v/v in PBS). Thereafter, bacteria were washed three times with PBS followed by incubation with Alexa Fluor 488 conjugates anti-mouse antibody against IgM (Life technologies, Eugene, OR, USA; A21042) and IgG (Life technologies, Eugene, OR, USA; A11029) at a dilution of 1:1000 (v/v in 1% HSA) for 3h at RT. After washing with PBS, bacteria were stained with 2µg/µL of 4’,6-diamidino-2-phenylindole (DAPI) and mounted in ProLong Diamond Antifade (Life Technologies, Eugene, OR, USA; P36961). Image acquisition was performed using a Leica confocal microscope (Leica, Wetzlar, Germany) with 63X oil immersion objective. Representative pictures were assembled in Adobe Illustrator and fluorescence was slightly enhanced using Adobe Photoshop CS6 (Adobe System Incorporated, California, USA).

### Tick infestation

Unfed nymphs were obtained from the colonies of UMR-BIPAR, Maisons-Alfort, France. Each mouse was infested with twenty *I. ricinus* nymphs on study day 40. Ticks were placed within EVA-foam (Cosplay Shop, Brugge, Belgium) capsules glued on the back of the animals as previously described (17). Unfed and fully-engorged nymphs were used for DNA extraction.

### DNA Extraction and 16S rRNA sequencing

Before DNA extraction, nymphs were washed two times in miliQ sterile water and one time in 70% ethanol. Ticks were pooled (5 ticks per pool) and crushed with glass beads using a Precellys24 Dual homogenizer (Bertin Technologies, Paris, France) at 5500× g for 20 s. Genomic DNA was extracted using a Nucleospin tissue DNA extraction Kit (Macherey-Nagel, Hoerdt, France). Each DNA sample was eluted in 100 µl of sterile water and DNA sequencing was commissioned to Novogene facility (London, UK) for amplicon sequencing of the bacterial 16S rRNA gene. A single lane of Illumina MiSeq system was used to generate 251-base paired-end reads from the V4 variable region of the 16S gene using barcoded universal primers (515F/806R). The raw 16S rRNA sequences were deposited at the SRA repository (Bioproject No. PRJNA725498, SRA accession No. SUB9549819). Four extraction reagent controls were set in which the different DNA extraction steps were performed under the same conditions as for the samples, but using water as template. DNA amplification was then performed on the extraction control in the same conditions as for any other sample.

### 16S rRNA sequences processing

The analysis of 16S rRNA sequences was performed using QIIME 2 pipeline (v. 2019.1) (18). The sequences in the fastq files were denoised and merged using the DADA2 software (19) as implemented in QIIME 2. The amplicon sequence variants (ASVs) were aligned with q2-alignment of MAFFT (20) and used to construct a phylogeny with q2-phylogeny of FastTree 2 (21). Taxonomy was assigned to ASVs using a classify-sklearn naïve Bayes taxonomic classifier (22) based on SILVA database (release 132) (23). Only the target sequence fragments were used in the classifier (i.e., classifier trained with the primers) (24,25). Taxa that persisted across serial fractions of the samples using QIIME 2 plugin feature-table (core-features) were considered ubiquitous (18).

### Bacterial co-occurrence networks, identification of keystone taxa and attack tolerance test

Co-occurrence networks were inferred for each dataset, based on taxonomic profiles, collapsed at the genus level. Correlation matrices were calculated using the SparCC method (26), implemented in the R environment. The topological parameters, i.e., the number of nodes and edges, weighted degree, centrality metrics and the hub-score of each node, the diameter of the network, modularity, and clustering coefficient were calculated for each network. Network calculations and visualizations were prepared with the software Gephi 0.9.2 (27). Three criteria were used to identify keystone nodes within the networks as previously described (16): (*i*) high eigenvector centrality values, (*ii*) ubiquitousness, and (*iii*) the combination of high relative abundance and eigenvector centrality values. The resistance of these taxonomic networks to taxa removal (i.e., attack tolerance) was tested on these taxonomic networks. The purpose was to measure their resistance to the systematic removal of nodes, either by a random attack with 100 iterations, or by a directed attack, removing the nodes according to its value of betweenness centrality (the highest, the first). The analysis of the network resistance was done with the package NetSwan for R (28).

### Prediction of functional traits in the tick microbiome

The 16S rRNA amplicon sequences from each data set were used to predict the metabolic profiling of each sample. PICRUSt2 (29) was used to predict the metagenomes from 16S rRNA amplicon sequences. Briefly, the AVSs were placed into a reference tree (NSTI cut-off value of 2) contained 20,000 full 16S rRNA sequences from prokaryotic genomes, which is then used to predict individual gene family copy numbers for each AVS. The predictions are based on Kyoto Encyclopedia of Genes and Genomes (KEGG) orthologs (KO) (30). The output of these analyses included pathways and EC (Enzyme Commission number) profiling; the pathways were constructed based on the MetaCyc database (31)

### Statistical analysis

Differences in relative antibody levels (i.e., OD) between groups of immunized mice in the different time points were compared using two-way ANOVA with Bonferroni multiple comparison tests applied for individual comparisons. Microbial diversity analyses were carried out on rarefied ASV tables, calculated using the q2-diversity plugins. The alpha diversity (richness and evenness) was explored using Faith’s phylogenetic alpha diversity index (32) and Pielou’s evenness index (33). Differences in α-diversity metric between groups were assessed using Kruskal-Wallis test (alpha= 0.05). Bacterial β-diversity was assessed using the Bray Curtis dissimilarity (34), and compared between groups using the PERMANOVA test. The differential abundant taxa and functional feature (KO genes and pathways) were explored between bacteria- and PBS-immunized mice, the differential features were detected by comparing the log2 fold change (LFC) using the Wald test as implemented in the compositional data analysis method DESeq2 (35).

Correlations between tick microbiota bacteria abundance and mice Abs levels were calculated with the ANOVA-Like Differential Expression (ALDEx2, v. 1.22.0) correlation analysis (aldex.cor function) as implemented in R (v. 4.0.3). Unpaired non-parametric Mann-Whitney U test was used to compare the tick parameters (i.e., time to complete feeding, the weight of engorged ticks and tick mortality) between groups. Two-way ANOVA and Mann-Whitney U test analyses were performed in the GraphPad 5 Prism software (GraphPad Software Inc., San Diego, CA, USA). Differences were considered significant when *p* < 0.05.

## Results

### Vaccination with keystone bacteria induces increased engorgement and reduced bacterial diversity in *Ixodes ricinus*

No mortality or adverse events were observed in mice immunized with *E. coli* or *L. mesenteroides*. Following the immunization protocol, each mouse was infested with 20 *I. ricinus* nymphs. Time to complete feeding, the weight of engorged ticks and tick mortality were recorded and compared between immunized and control groups. A significant increase (Mann-Whitney U test, *p* = 0.03) in weight was recorded in nymphs that fed on *E. coli*-immunized mice compared with the mice of the control group (Figure 1A). This was not the case for ticks engorged on *L. mesenteroides*-immunized mice (Mann-Whitney U test, *p* > 0.05). There were no significant differences (Mann-Whitney U test, *p* > 0.05) in the total number of ticks that dropped naturally (Figure 1B) or mortality of ticks that fed on *E. coli*-immunized, *L. mesenteroides*-immunized, or control mock-immunized mice (Figure 1C).

**Figure 1.**
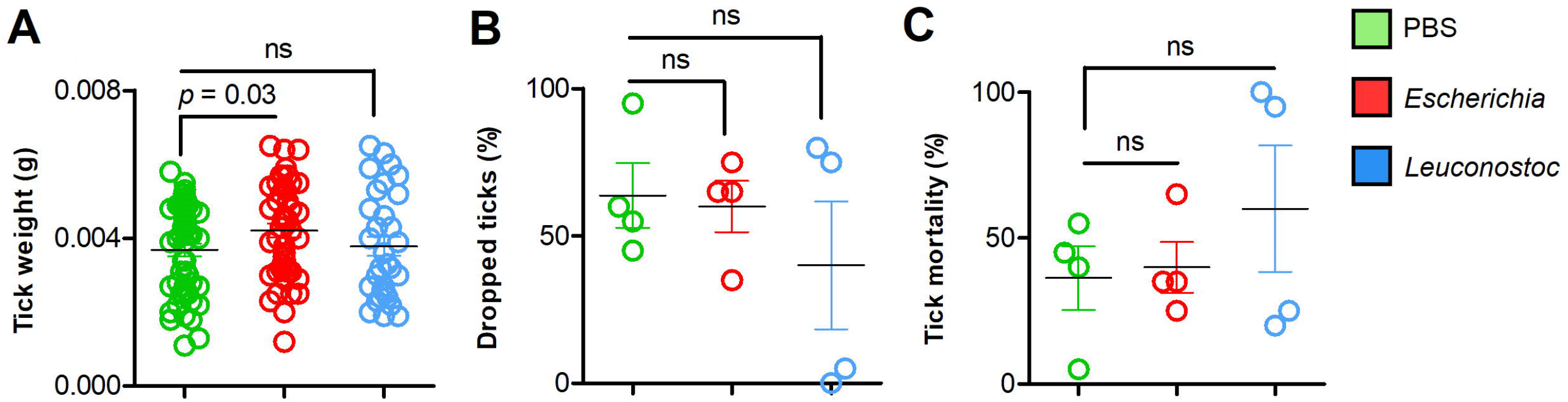
Performance of *I. ricinus* nymphs feeding on mice vaccinated with live *E. coli* or *L. mesenteroides*. (A) The weight of individual engorged ticks was measured and compared between groups. (B) The percentage of ticks that engorged and dropped was calculated and compared between groups. (C) The percentage of dead ticks was calculated and tick mortality (%) was compared between groups. Means and standard deviation values are displayed. The parameters were compared between groups by the Mann-Whitney U test. (ns-not significant; n = 12 mice (*n* = 20 ticks per mouse)

The impact of anti-microbiota vaccines on the diversity, composition and abundance of tick microbiota bacteria was assessed after 16S rRNA amplicon sequencing of DNA extracted from unfed *I. ricinus* nymphs or after engorgement on *E. coli*-immunized, *L. mesenteroides*-immunized, or mock-immunized mice. Vaccination with the keystone bacterium *E. coli* decreased the bacterial diversity associated with the tick microbiota (H = 8.6, *p* = 0.03, Figure 2A), but had no significant impact (H = 5.8; *p* = 0.12) on the species evenness (Figure 2B). Conversely, vaccination with the non-keystone bacterium *L. mesenteroides* had no impact on bacterial diversity (Figure 2A), but decreased significantly the species evenness, compared to unfed nymph (Figure 2B). Overall, the comparison of the diversity indexes of unfed and fed ticks revealed that anti-microbiota vaccination interferes with the normal dynamics of tick microbiota, regardless of the keystoneness of the bacteria used in the vaccine formulation. Accordingly, a Principal Coordinates Analysis (PCoA) shows that the profiles of both groups of ticks that fed on bacteria-immunized mice are very similar compared to mock-immunized or unfed ticks (*F*= 5.30; *p* = 0.00, Figure 2C).

**Figure 2.**
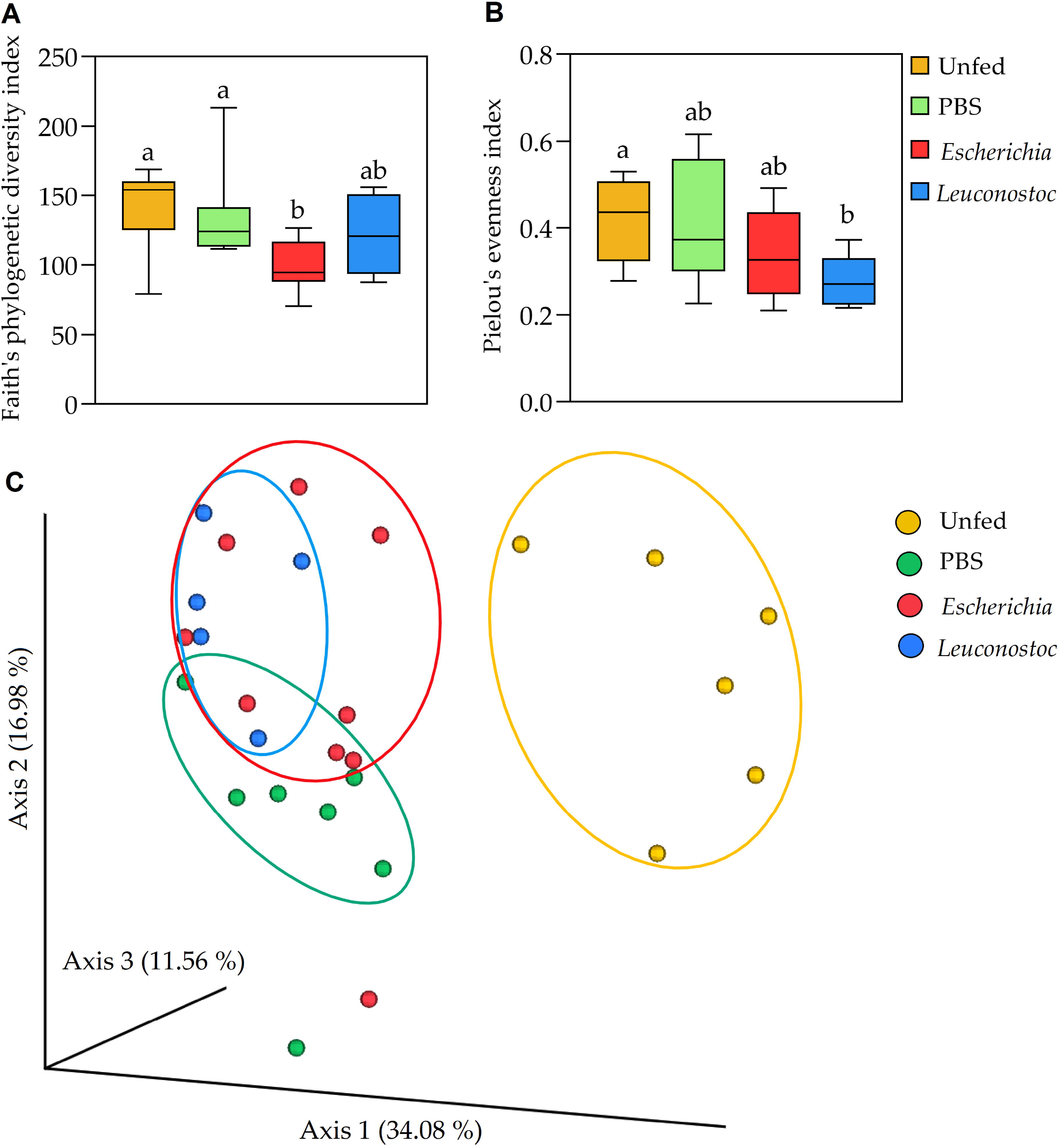
Impact of anti-microbiota vaccines on tick microbial diversity and evenness. (A) Faith’s phylogenetic diversity and (B) Pielou’s evenness indexes were used to calculated the microbial richness and evenness, respectively, of the bacterial communities of unfed ticks and ticks fed on mock-immunized (green, PBS), *E. coli*-immunized (red) and *L. mesenteroides*-immunized (blue) mice. (C) First axis PCoA plot showing the variance of taxonomical profile (Bry Curtis distance) at the level of genera between samples according to tick feeding status and feeding on immunized mice (PERMANOVA, *p* = 0.00), arbitrary ellipses were drawn to facilitate the interpretation of the figure.

### Increase of bacteria-specific Abs was associated with reduced abundance of the keystone taxon *Escherichia*-*Shigella*

Taxa composition and abundance analysis showed significant changes in the abundance of several bacterial genera in ticks collected from mock-immunized mice compared with unfed ticks (Figure 3A). There were significant changes in taxa abundance in the ticks engorged on either *L. mesenteroides*-immunized (Figure 3B), or *E. coli*-immunized (Figure 3C) mice compared to the mock-immunized group. The taxa with significant changes in abundance, measured as centered log ratio (clr), are displayed in figures 3D and 3E for *L. mesenteroides*-immunized and *E. coli*-immunized mice, respectively. Notably, the abundance of *Escherichia*-*Shigella*, but not of *Leuconostoc*, was significantly reduced in ticks that fed on *E. coli*-immunized mice compared with the control group (Figure 3E). In contrast, the abundance of *Escherichia*-*Shigella* or *Leuconostoc* was not significantly affected in ticks that fed on *L. mesenteroides*-immunized mice (Figure 3D).

**Figure 3.**
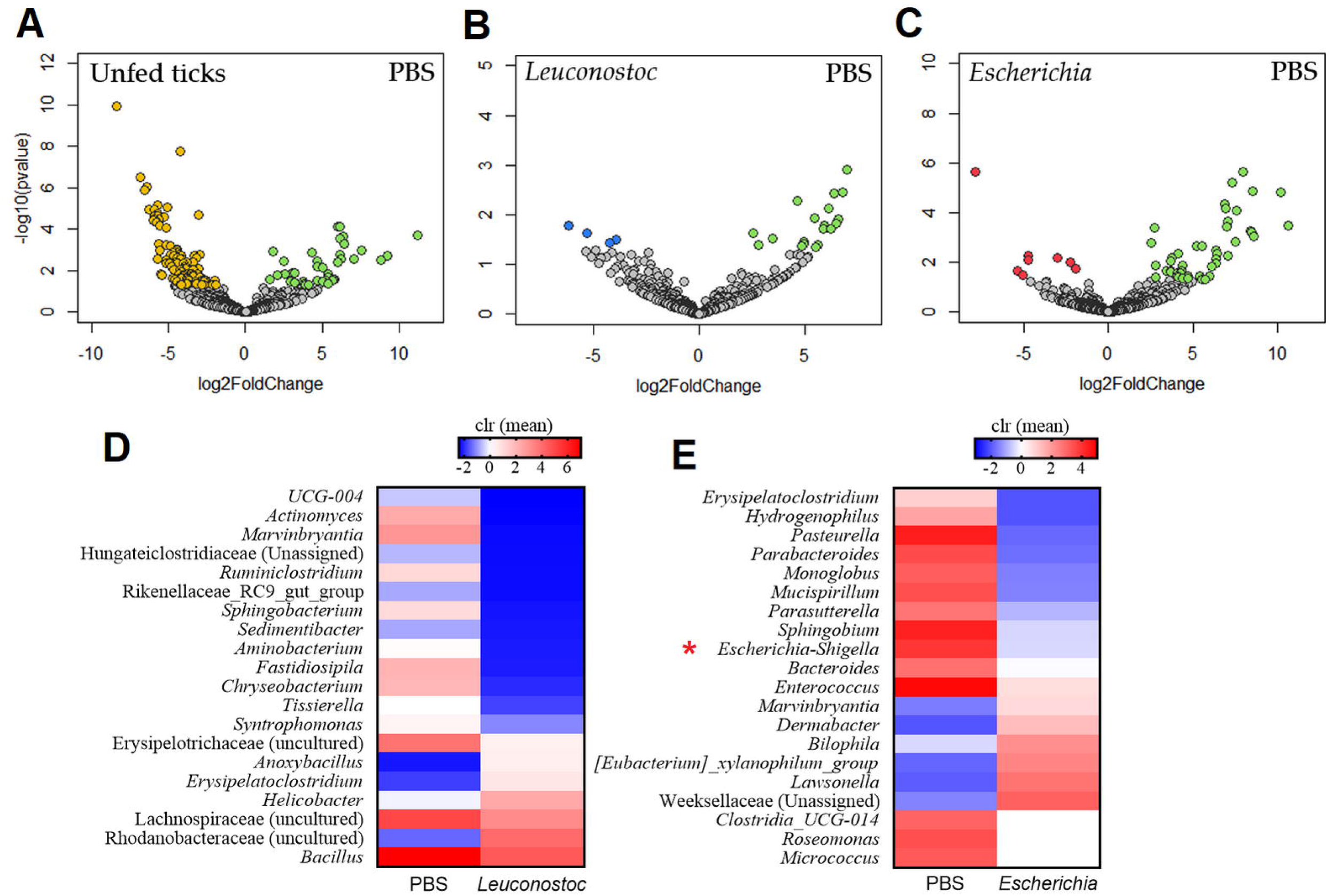
Impact of anti-microbiota vaccines on the taxonomic profiles of tick microbiome. Volcano plot showing differential bacterial abundance in ticks of the different groups: (A) unfed ticks vs. ticks fed on mock-immunized mice (PBS), (B) ticks fed on mock-immunized vs. *L. mesenteroides*-immunized mice, and (C) ticks fed on mock-immunized vs. *E. coli*-immunized mice. The yellow (unfed ticks), blue (ticks fed on *L. mesenteroides*-immunized mice), and red (ticks fed on *E. coli*-immunized mice) dots indicate taxa that displayed both large magnitude fold-changes and high statistical significance favoring disturbed or control group (green dots), while the gray dots are considered as not significantly different between groups. The relative abundance (calculated as clr transformed values) of the 20 top bacterial taxa with the highest significant differences on ticks fed on mock-immunized vs. *L. mesenteroides*-immunized mice (D) and on ticks fed on mock-immunized vs. *E. coli*-immunized mice (E), as detected by the DeSeq2 algorithm (Wald test, *p* < 0.001).

Immunization with live *E. coli* induced a significant increase in IgM and IgG specific to *E. coli* proteins (Figure 4A). Strong and specific immune reaction of mice IgM against *E. coli* on d30 was confirmed by immunofluorescence (Supplementary Figure S1A). The reaction of anti-*E. coli* IgG from *E. coli*-immunized mice against *E. coli* was also confirmed by immunofluorescence (Supplementary Figure S1B). However, immunization with live *L. mesenteroides* elicited only marginal levels of IgM on d30 that dropped by d46 (Figure 4B), suggesting that in contrast to *E. coli, L. mesenteroides* is poorly immunogenic as a live vaccine. Consistent with the low IgM levels raised against the bacterium *L. mesenteroides* on d30 (Figure 4B), we observed a modest recognition of *L. mesenteroides* by mice IgM immunized with the gram-negative bacteria (Supplementary Figure S1C). No reaction was observed for when anti-*L. mesenteroides* IgG detection was used (Supplementary Figure S1D), which is consistent with the low levels of anti-*L. mesenteroides* IgG detected by ELISA (Figure 4B).

**Figure 4.**
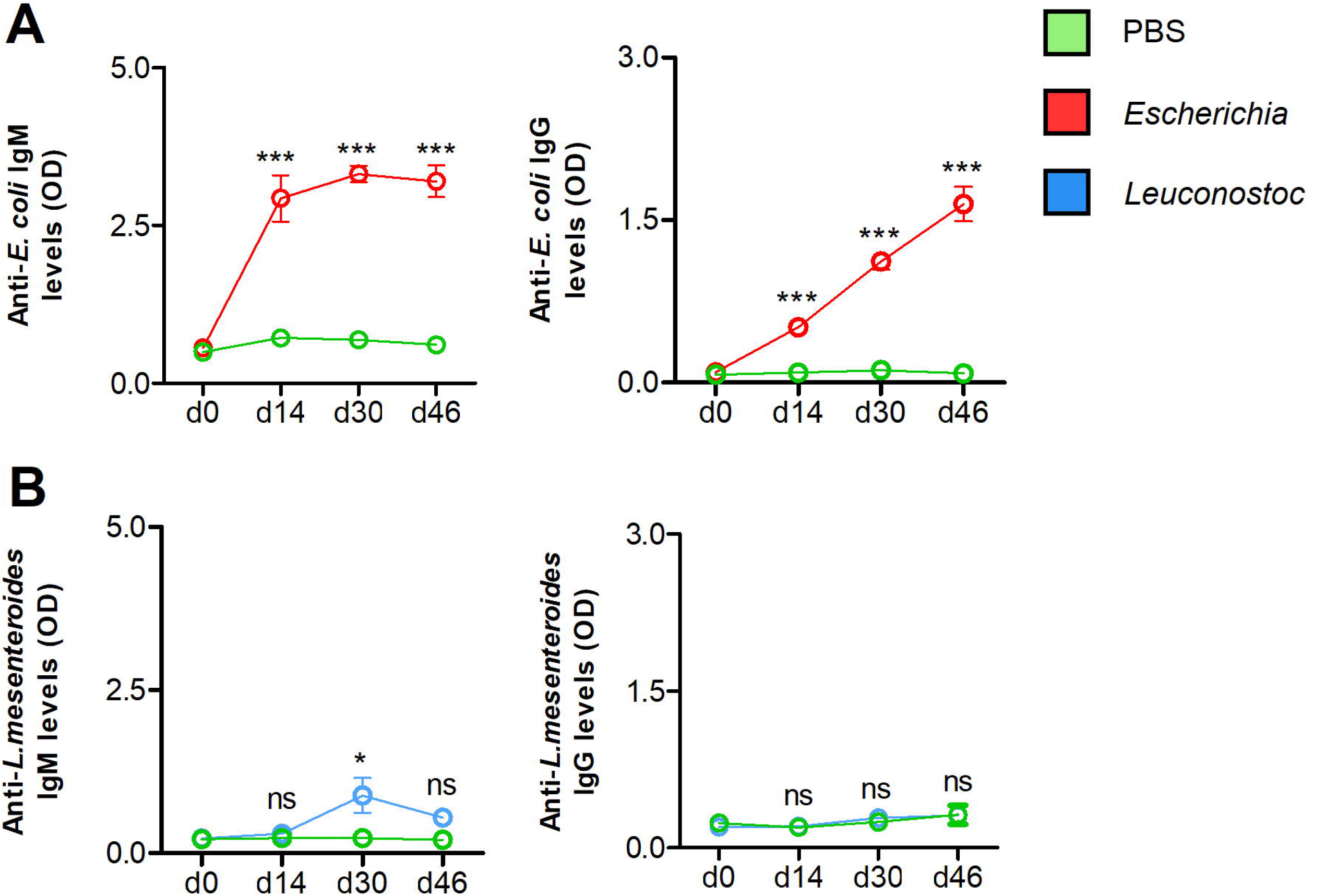
Antibody response of mice vaccinated with live *E. coli* or *L. mesenteroides*. The levels of IgM and IgG specific to (A) *E. coli* and (A) *L. mesenteroides* proteins were measured by semi-quantitative ELISA in sera of mice immunized with *E. coli* (red) and *L. mesenteroides* (blue), respectively, at different time points (days, d), d0, d14, d30 and d46. Antibody levels of bacteria-immunized mice were compared with those of mock-immunized (green, PBS) mice. Means and standard error values are shown. Results were compared by two-way ANOVA with Bonferroni test applied for comparisons between control and immunized mice. (* *p* < 0.05, *** *p* < 0.0001; ns-not significant; 1 experiment, n = 12 mice and three technical replicates per sample.

No significant cross-reaction to *E. coli* proteins was detected in the IgM and IgG fractions of the sera of mice immunized with the live vaccine containing *L. mesenteroides*. A weak increase in anti-*E. coli* IgM on d30 and d46, and no increase in anti-*E. coli* IgG were observed in *L. mesenteroides*-immunized mice (Supplementary Figure S2A). However, anti-*L. mesenteroides* IgG increased on d46, and IgM specific to proteins of this bacterium increased in response to *E. coli* vaccination (Supplementary Figure S2B).

To test a possible association between the increase in *E. coli*-specific IgM and IgG after vaccination and the decrease in the abundance of *Escherichia*-*Shigella*, we performed an ALDEx2 correlation analysis between the abundance of all bacterial taxa at genus level and Abs levels. Significant negative correlations were found between the levels of anti-*E. coli* IgM (*r*_s_ = −0.60, *p* = 0.01) and IgG (*r*_s_ = −0.57, *p* = 0.02) in mice sera and the abundance of *Escherichia*-*Shigella* in the ticks. A negative correlation was also found between anti-*E. coli* IgM (*r*_s_ = −0.57, *p* = 0.02) and IgG (*r*_s_ = −0.64, *p* = 0.01) levels and one bacteria genus (0.18%, total 533 taxa), *Parabacteroides*, family Porphyromonadaceae. A positive correlation was found between the genus *Lawsonella*, Order Corynebacteriales, and anti-*E. coli* IgM (*r*_s_ = 0.65, *p* = 0.01) and IgG (*r*_s_ = 0.68, *p* = 0.008) levels. No taxa abundance was found to correlate with the levels of both anti-*E. coli* IgM and IgG in the *L. mesenteroides*-immunized mice. In addition, no statistically significant correlations were found between the anti-*L. mesenteroides* IgM and IgG levels and the abundance of *Leuconostoc*, or any other taxa identified in the tick microbiome. Taken together, these results showed that anti-*E. coli* immunization of mice reduces the *Escherichia*-*Shigella* abundance within the tick microbiome in a taxon-specific manner.

### Anti-tick microbiota vaccine reshapes the hierarchy of nodes in co-occurrence networks

The taxonomic profiles generated were used to build a co-occurrence network and the eigenvector centrality values were computed for each node. Bacteria with a high eigenvector centrality value (> 0.90) were considered as keystone bacteria in the microbiota of *I. ricinus*. The impact of live bacteria immunization on the structure of the tick microbial communities was then quantified and visualized using co-occurrence networks. In accordance with their classification as non-keystone and keystone taxa in the networks of ticks that fed on mock-immunized mice (Supplementary Figure S3A), *Leuconostoc* and *Escherichia*-*Shigella* were poorly (Figure 5A) and highly (Figure 5B) interconnected to other taxa, respectively. Visual inspection of local connectedness around *Leuconostoc* and *Escherichia*-*Shigella* reveals that tick feeding on mice immunized with these bacteria exhibited an increase (Figure 5C) and decrease (Figure 5D) in the number of co-occurring taxa. Notably, the eigenvector centrality value of *Leuconostoc* was very similar in the networks inferred from ticks fed on *L. mesenteroides*-immunized (eigenvector 0.11) and mock-immunized (eigenvector 0.12) mice, while the eigenvector value of *Escherichia*-*Shigella* decreased 95 times in the network of ticks fed on *E. coli*-immunized (eigenvector 0.01) compared with those fed on mock-immunized mice (eigenvector 0.95). Visual (Supplementary Figure S3) and numerical (Table 1) comparison of network parameters show that, in addition to the local connectedness effect, anti-microbiota vaccination had a large impact on the structure of the microbial community of ticks. For instance, the number of edges decreased in the co-occurring networks of ticks that fed on *E. coli*-immunized compared to the control group (Table 1) and increased in the networks of ticks that fed on *L. mesenteroides*-immunized mice.

**Table 1.**
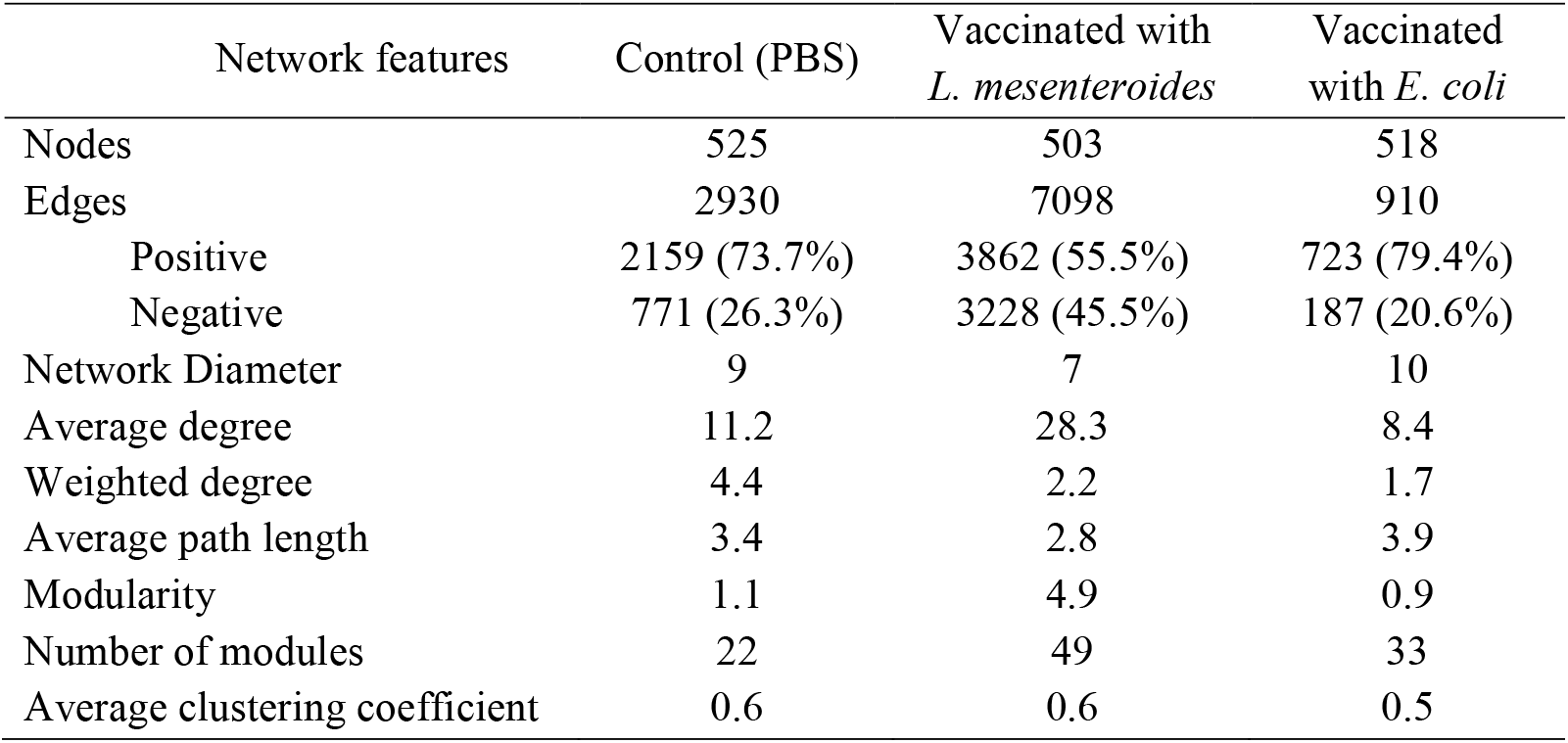
Topological parameters of co-occurrence networks.

| Network features | Control (PBS) | Vaccinated with *L. mesenteroides* | Vaccinated with *E. coli* |
| --- | --- | --- | --- |
| Nodes | 525 | 503 | 518 |
| Edges | 2930 | 7098 | 910 |
| Positive | 2159 (73.7%) | 3862 (55.5%) | 723 (79.4%) |
| Negative | 771 (26.3%) | 3228 (45.5%) | 187 (20.6%) |
| Network Diameter | 9 | 7 | 10 |
| Average degree | 11.2 | 28.3 | 8.4 |
| Weighted degree | 4.4 | 2.2 | 1.7 |
| Average path length | 3.4 | 2.8 | 3.9 |
| Modularity | 1.1 | 4.9 | 0.9 |
| Number of modules | 22 | 49 | 33 |
| Average clustering coefficient | 0.6 | 0.6 | 0.5 |

**Figure 5.**
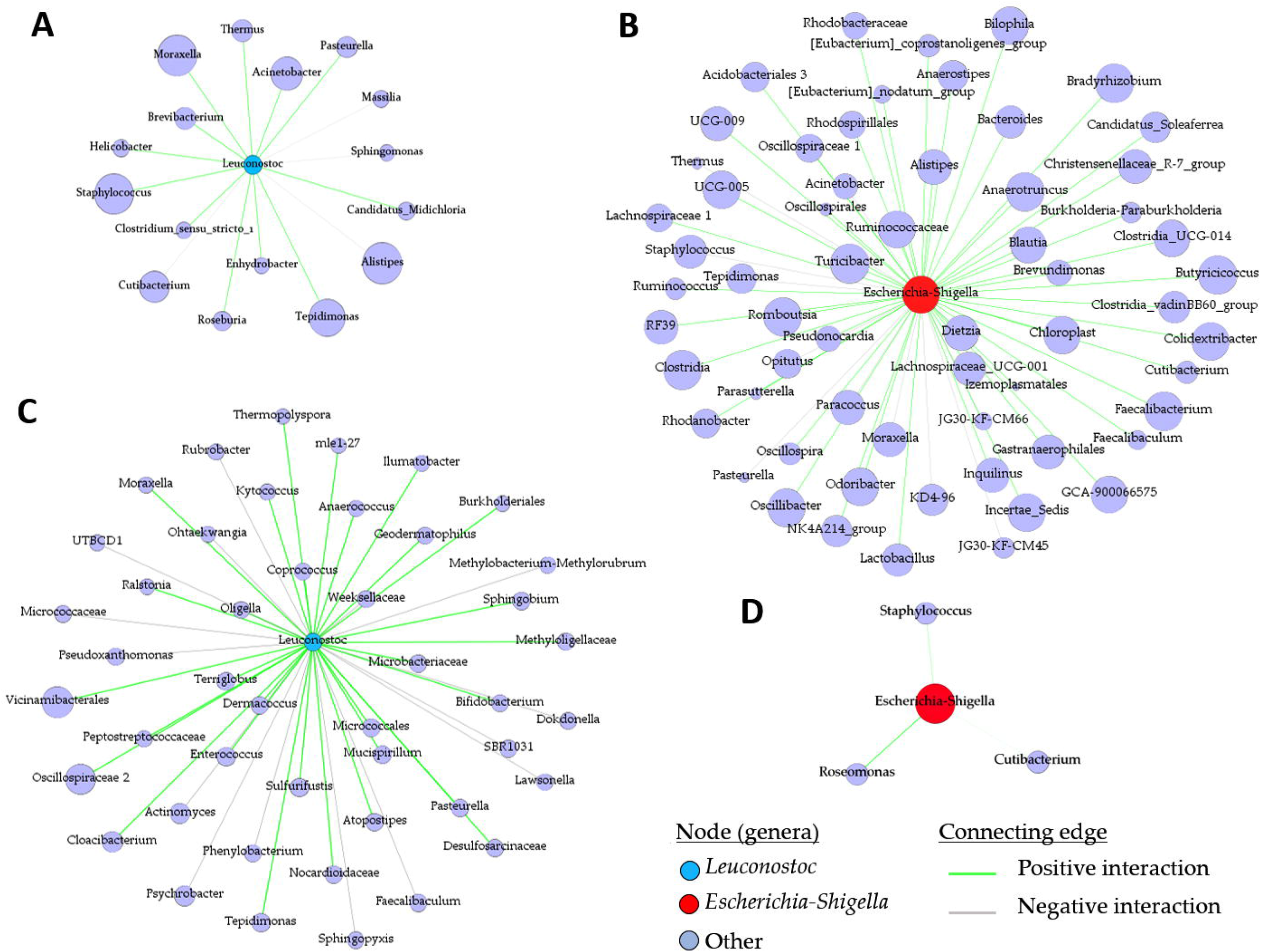
Local connectivity of *Leuconostoc* and *Escherichia*-*Shigella* in the co-occurrence networks. The nodes/taxa linked to *Leuconostoc* (cyan node, A, C) and *Escherichia*-*Shigella* (red node, B, D) were identified in the bacterial co-occurrence networks of ticks fed on mock-immunized (A, B), *L. mesenteroides*-immunized (C) and *E. coli*-immunized (D) mice. Connecting edges with positive and negative interactions were also identified.

Taxonomic networks were tested for attack tolerance. In this analysis, the resistance of the networks to random or directed, removal of nodes was measured and the proportion of taxa removal needed to reach a loss in connectivity of 0.90 was recorded for each network. A proportion of 0.58, 0.55 and 0.66 randomly removed nodes produce a 0.90 connectivity loss in the networks of ticks from the control (Figure 6A), *E. coli*-immunized (Figure 6B) and *L. mesenteroides*-immunized (Figure 6C) mice, respectively. The same loss in connectivity (i.e., 0.90) was observed when a smaller proportion (i.e., 0.23 for the control group, 0.14 for *E. coli* group and 0.53 for the *L. mesenteroides* group) of highly central nodes were removed first from each network (Figure 6). Thus, immune targeting of the keystone taxon *Escherichia*-*Shigella* decreased attack tolerance in bacterial co-occurrence network. The random and directed removal curves within the network of ticks from *L. mesenteroides*-immunized mice revealed high similarity, which was not the case in the other two networks. This suggests an unstructured hierarchy of nodes in the co-occurrence network of ticks from *L. mesenteroides*-immunized mice, and a reshaping in the hierarchy of nodes in the co-occurrence network of ticks from *E. coli*-immunized mice compared to the control group.

**Figure 6.**
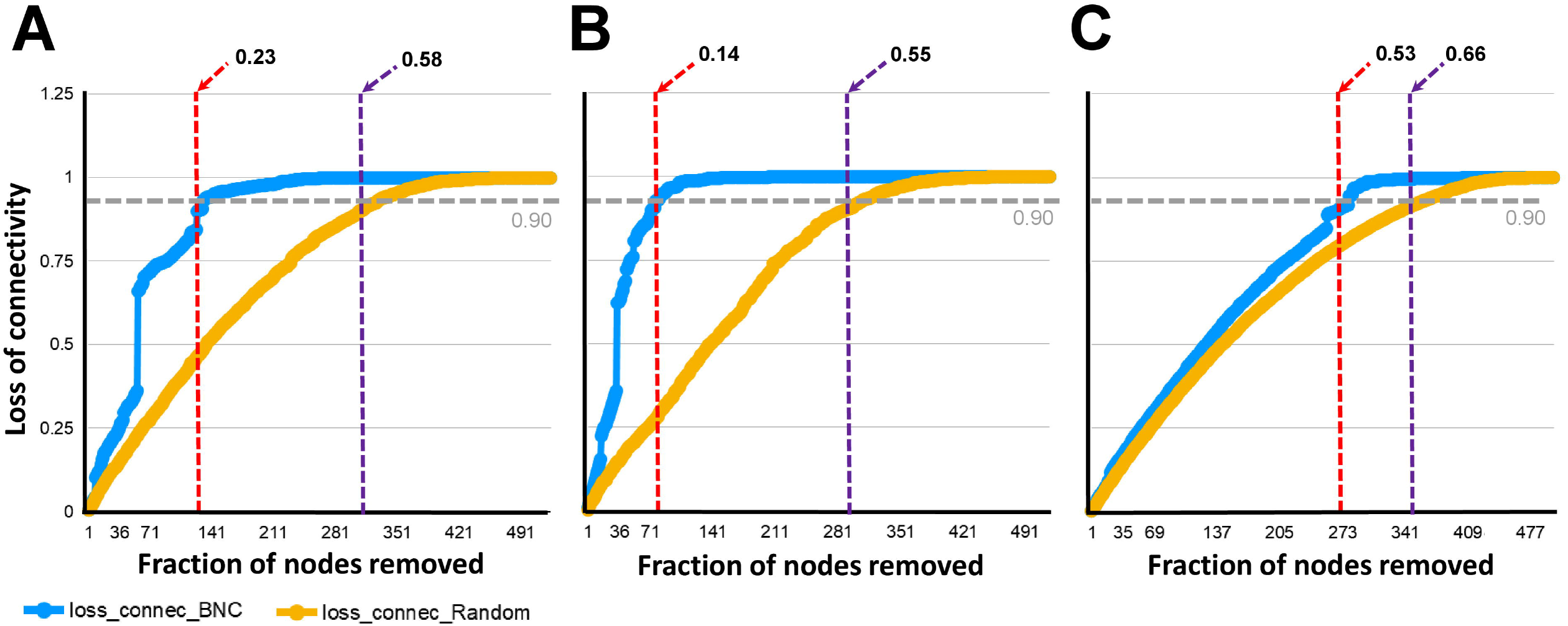
Network tolerance to node removal. The resistance of the networks to directed (blue line), or random (orange line), removal of nodes was measured. The proportion of directly (red dashed line), or randomly (violet dashed line) removed nodes that made the network losing 0.90 of connectivity was recorded in the bacterial co-occurrence networks of ticks fed on mock-immunized (A), *L. mesenteroides*-immunized (B) and *E. coli*-immunized (C). Loss of connectivity values range between 0 (maximum of connectivity between nodes) and 1 (total disconnection between nodes) for any given network.

### Community disturbance by anti-microbiota vaccine changes the abundance of putative metabolic pathways in the tick microbiome

We analyzed microbial putative functions displayed by the microbiome of unfed ticks and those that fed on *L. mesenteroides*-immunized and *E. coli*-immunized mice by PICRUSt2 and compared them with the bacterial community predicted functions present in the ticks that fed on mice immunized with the mock vaccine (Figure 7). Blood feeding on control mice induced fold changes in the relative abundance of several putative genes (KO) in *I. ricinus* microbiome compared to unfed ticks (Figure 7A). Major changes in the predicted gene profiles were also observed in the microbiome of tick from *L. mesenteroides*-immunized (Figure 7B) and *E. coli*-immunized (Figure 7C) mice. Putative pathway analysis revealed that some of the functional pathways were conserved and others were specific to each group (Figure 8), when considering those with significant log2fold change above 1 in their abundance (Figure 7A). A total of 115 pathways had differential abundance (81 and 34 with decreased and increased abundance, respectively, Log2fold change > 1, *p* < 0.05) in the ticks that fed on mock-immunized mice compared to unfed ticks (Supplementary Table S1). Thirteen pathways had a log2fold change lower than −1 in the functional prediction of the microbiome of ticks fed on *L. mesenteroides*-immunized mice, compared to the control group of ticks fed on mock-immunized mice (Supplementary Table S2). Only one of them, tetrahydromethanopterin biosynthesis (Log2fold change = −5.5, Kruskal-Wallis test, *p* = 0.02), changed exclusively in the ticks of the *L. mesenteroides*-immunized group. Fourteen pathways had a log2fold change lower than −1 in the functional prediction of the microbiome of ticks fed on *E. coli*-immunized mice, compared to the control group (Supplementary Table S3). A significant decrease in the relative abundance of only the L-lysine fermentation to acetate and butanoate (Log2fold change = −1.6, Kruskal-Wallis test, *p* = 0.008) pathway was found exclusively in ticks fed on *E. coli*-immunized mice. The relative abundance of only a reduced number of pathways (i.e., methanogenesis from acetate, super pathway of glycerol degradation to 1,3-propanediol, superpathway of (Kdo)2-lipid A biosynthesis, and CMP-legionaminate biosynthesis I) changed significantly in the three groups of fed ticks and they may represent functional changes induced by blood feeding in the bacterial communities independent of the treatment.

**Figure 7.**
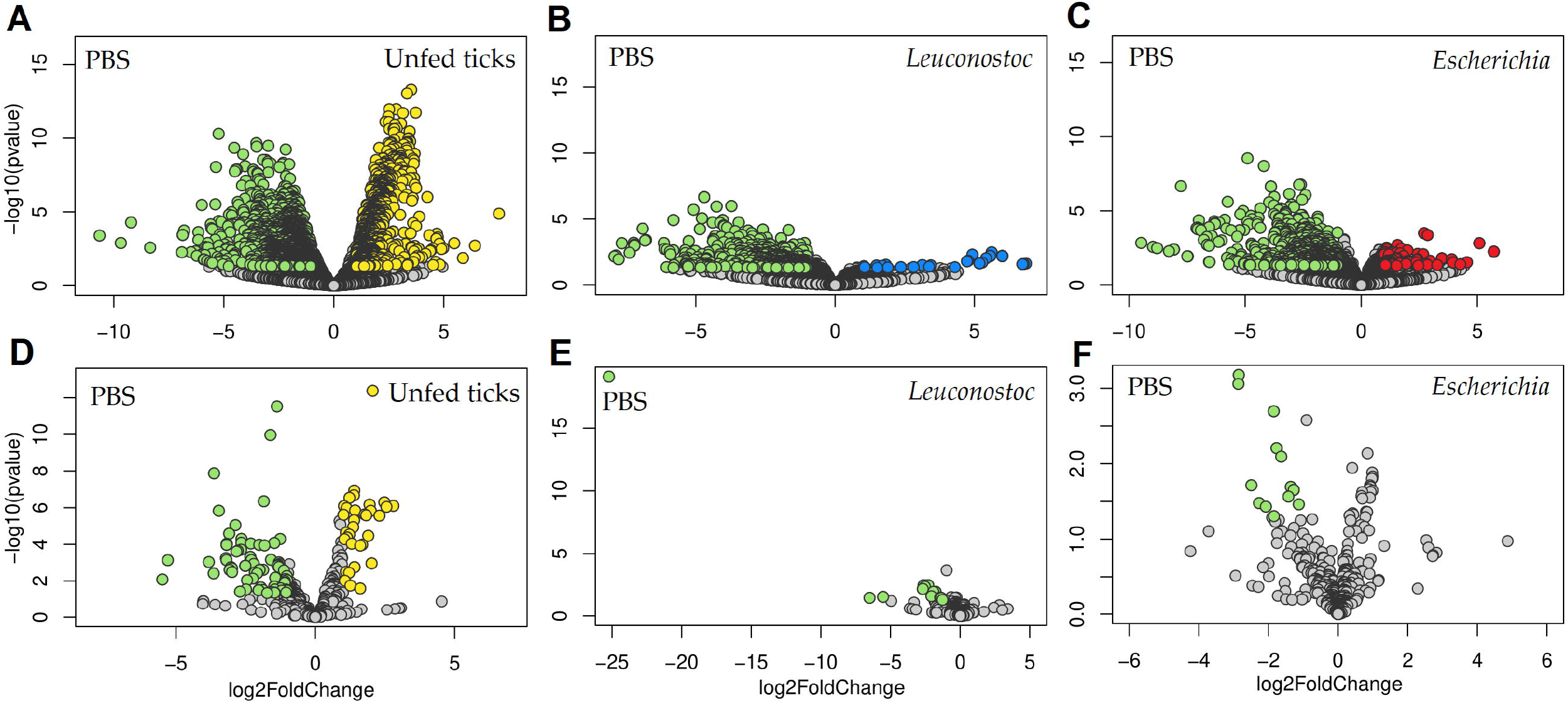
Impact of anti-microbiota vaccines on the predicted functional profiles of tick microbiome. Volcano plot showing differential enzyme (A-C) and pathway (D-F) abundance in ticks of the different groups: (A and D) unfed ticks vs. ticks fed on mock-immunized mice (PBS), (B and E) ticks fed on mock-immunized vs. *L. mesenteroides*-immunized mice, and (C and F) ticks fed on mock-immunized vs. *E. coli*-immunized mice. The yellow (unfed ticks), blue (ticks fed on *L. mesenteroides*-immunized mice), and red (ticks fed on *E. coli*-immunized mice) dots indicate enzyme (A-C) and pathway (D-F) that displayed both large magnitude fold-changes and high statistical significance favoring disturbed or control PBS group (green dots), while the gray dots are considered as not significant. Differential features were detected by the DeSeq2 algorithm (Wald test, *p* < 0.05).

**Figure 8.**
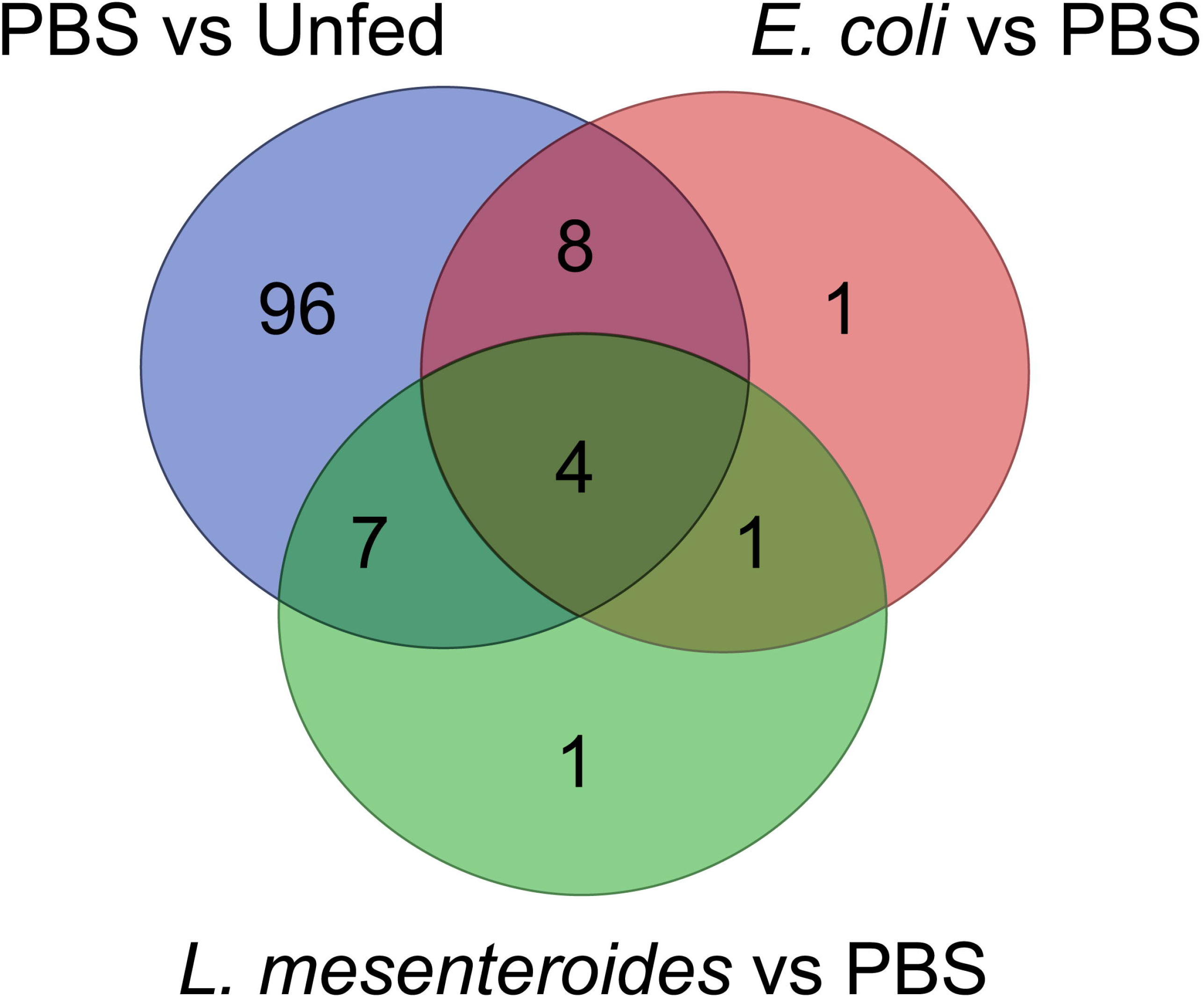
Differential predicted pathways influenced by feeding and anti-microbiota vaccines. Venn diagram showing the common and different predicted bacterial pathways modulated by feeding (unfed ticks vs. ticks fed on mock-immunized mice), and anti-microbiota vaccination in ticks fed on mock-immunized vs. *L. mesenteroides*-immunized mice, and ticks fed on mock-immunized vs. *E. coli*-immunized mice. Only pathways with statistically significant log2 fold changes of absolute value cutoff of 1 were considered.

## Discussion

Several studies have shown that the tick microbiome is a gate to access tick physiology and vector competence (1,36). Specifically, a reduced bacterial load has been associated with decreased reproductive fitness after antibiotics treatment in ticks (3,36-40). However, considering that antibiotics can target several bacteria simultaneously, studies using broad-spectrum antimicrobial compounds make impossible to stablish causal links between specific taxa and their role on tick physiology. Recently, we showed for the first time that anti-tick microbiota vaccines impact tick performance during feeding (16). Considering that host Abs and complement acquired during tick feeding not only retain their immune functions, but also access several tick tissues (40-45), we hypothesized that anti-tick microbiota vaccines can be used as a precision microbiology tool to target selected taxa in the tick microbiome. Here we show that immunization with a live *E. coli* vaccine elicited bacterial-specific Abs of the isotypes IgM and IgG. Furthermore, immune targeting of the keystone taxon *Escherichia*-*Shigella* using a live *E. coli* vaccine reduced the relative abundance of *Escherichia*-*Shigella* in engorged ticks, suggesting that host Abs can bind and promote killing bacteria within tick microbiota. Antibodies induced against particular tick proteins can also react with the corresponding tick protein within tick tissues (43,46). For example, host Abs against the glycoprotein Bm86, predominantly located in the membrane of tick gut cells (46), modulate tick proteome (47) and bind to the surface of epithelial cells in the tick intestine causing cell lysis (43). Therefore, it can be presumed that when ingested during blood feeding, the Abs produced against selected tick microbiota bacteria could interfere with the physiological functionality of microbes within the ticks.

Targeting the keystone bacteria *Escherichia*-*Shigella* with host Abs also reduced its keystoneness, which was associated with a global modulation of the microbial community structure. The resulting community had a reduced alpha diversity, and changes in the taxonomic and predicted functional profiles. In addition, the co-occurring networks showed that *Escherichia*-*Shigella*-depleted communities had fewer nodes and the connectivity between them was weak and more susceptible to taxa extinction when compared with the control group of ticks that fed on mock-immunized mice. Removal of keystone species has strong disturbing effects, resulting in loss of microbiota biodiversity in different ecological settings (48-51). Depletion of keystone species could also result in microbial dysbiosis that impairs the integrity of the gut ecosystem, as seen in vertebrates (52,53). The impact of *E. coli* deletion has been tested experimentally on a synthetic consortium of 14 gut microbes (53). In particular, removal of *E. coli* resulted in the highest impact on biomass and growth rates, indicating major roles of this microorganism on a synthetic microbial consortium (53).

The bacterial community of ticks fed on *E. coli*-immunized mice had a significant decrease in the relative abundance of the predicted pathway L-lysine fermentation to acetate and butanoate. The relative abundance of other predicted pathways changed in response to feeding alone or feeding on *L. mesenteroides*-immunized mice. We consider the changes observed in the microbiome of ticks fed on *mesenteroides*-immunized mice as non-specific, or at least not dependent on a specific Abs response. This result suggests that bacterial community modulation by anti-microbiota vaccines could impact the functional profiles associated with the tick microbiome. Considering that lysine is an essential amino acid and that the tick genome does not encode for lysine synthesis enzymes (54), we hypothesized that the decrease of lysine degradation by tick microbiome might result in higher levels of free lysine available for tick metabolism. This could potentially explain the higher body weight of ticks fed on *E. coli*-immunized mice compared to mock-immunized and *L. mesenteroides*-immunized mice. The limitation here is that we used pathway prediction and did not perform metabolomics to test whether the availability of free lysine increases in the gut of ticks fed on *E. coli*-immunized mice. Further investigation is needed to examine the metabolite dynamics *in vivo* in response to bacterial community modulation by anti-*E. coli* IgM and IgG.

## Conclusions

In this study, we demonstrated that after immunization against the keystone microbiota taxon *E. coli*, the abundance of *Escherichia-shigella* in the tick microbiome negatively correlated with the level of host Abs specific to *E. coli* suggesting that anti-tick microbiota vaccine has the capacity to target a specific taxon through an immune response. We also showed that tick engorgement, microbiome bacterial diversity and microbial community structure can be disturbed by vaccination with *E. coli* highlighting the important role that keystone microbiota bacteria have in tick performance and microbiome. Specific-taxon changes in enzymes and pathways abundances observed with live bacteria vaccines suggest that the scope of anti-tick microbiota vaccine is not limited to the taxonomic modulation level, but it may also regulate the functions associated with the microbiome. In summary, targeting keystone bacteria of the tick microbiota by host Abs seems to be a suitable tool for the modulation of tick microbiome to study the role of a specific taxon in tick physiology. Anti-tick microbiota vaccine can also be a powerful tool to evaluate the functional contribution of a specific taxon in tick microbiota on pathogen colonization and transmission. These results guide precise interventions for the control of tick infestations and pathogen infection/transmission.

## Acknowledgements

UMR BIPAR is supported by the French Government’s Investissement d’Avenir program, Laboratoire d’Excellence “Integrative Biology of Emerging Infectious Diseases” (grant no. ANR-10-LABX-62-IBEID). Alejandra Wu-Chuang is supported by Programa Nacional de Becas de Postgrado en el Exterior “Don Carlos Antonio López” (grant no. 205/2018).

## Author contributions

AC-C, DO, JM and LM-H conceived the study. LM-H, AW-C, JM and JB performed the experiments and acquired the data. DO, LM-H, SD-S, AE-P and AC-C analyzed the data. DO, AC-C, AW-C and AE-P prepared figures and supplementary materials. LGB-H, ET-M, NV and JdlF contributed reagents and other resources. AC-C, LS, LGB-H, and JdlF supervised the work. AC-C, LM-H, AW-C, AH and DO drafted the first version of the manuscript. All the authors made editorial contributions, revised and accepted the final version of the manuscript.

## Supplementary materials

**Supplementary Figure S1. Immunocytochemistry of *E. coli* and *L. mesenteroides* using sera of immunized mice**. Fixed *E. coli* (A,B) and *L. mesenteroides* (C,D) were stained with pooled sera of mice immunized with a live *E. coli* vaccine (*Escherichia*), live *L. mesenteroides* vaccine (*Leuconostoc*) or mock vaccine (PBS). Examples of positive reaction are displayed (white arrows in inserts). Alexa fluor 488 conjugated anti-mouse antibody specific to the isotypes IgM (A,C) and IgG (B,D) were used as a secondary antibody. Negative control staining (Control) was performed using only the secondary antibody. Blue color indicates the nuclei visualized by 4’,6-diamidino-2-phenylindole (DAPI). Images were obtained using 63X magnification and digital zoom. Scale bars are 2µm.

**Supplementary Figure S2. Antibody response of mice vaccinated with live *E. coli* or *L. mesenteroides***. The levels of IgM and IgG specific to (A) *E. coli* and (B) *L. mesenteroides* proteins were measured by semi-quantitative ELISA in sera of mice immunized with *L. mesenteroides* (blue) and *E. coli* (red), respectively, at different time points, d0, d14, d30 and d46. Antibody levels of bacteria-immunized mice were compared with those of mock-immunized (green, PBS) mice. Means and standard error values are shown. Results were compared by two-way ANOVA with Bonferroni test applied for comparisons between control and immunized mice. (* *p* < 0.05, ** *p* < 0.001, *** *p* < 0.0001; ns-not significant; 1 experiment, n = 12 mice and three technical replicates per sample.

**Supplementary Figure S3**. A schematic representation of the co-occurring microbial taxa in the microbiome of ticks fed on mock-immunized (A), *L. mesenteroides*-immunized (B) and *E. coli*-immunized (C) mice. Circles (nodes) are bacterial genera and edges the co-occurrence between taxa. Equal colors mean clusters of taxa that co-occur more frequently among them than with other taxa. The size of the circles is proportional to the eigencentrality of each taxon in the resulting network. The nodes *Escherichia*-*Shigella* (red) and *Leuconostoc* (cyan) were identified and labelled (lighting symbol).

**Supplementary Table S1**. Pathways with differential abundance in the ticks that fed on mock-immunized mice compared to unfed ticks.

**Supplementary Table S2**. Pathways with differential abundance in the ticks that fed on *L. mesenteroides*-immunized compared to mock-immunized mice.

**Supplementary Table S3**. Pathways with differential abundance in the ticks that fed on *E. coli*-immunized compared to mock-immunized mice.

