## Supplementary figures and images for "Anti-microbiota vaccines modulate the tick microbiome in a taxon-specific manner"

### Supplementary Figure S1

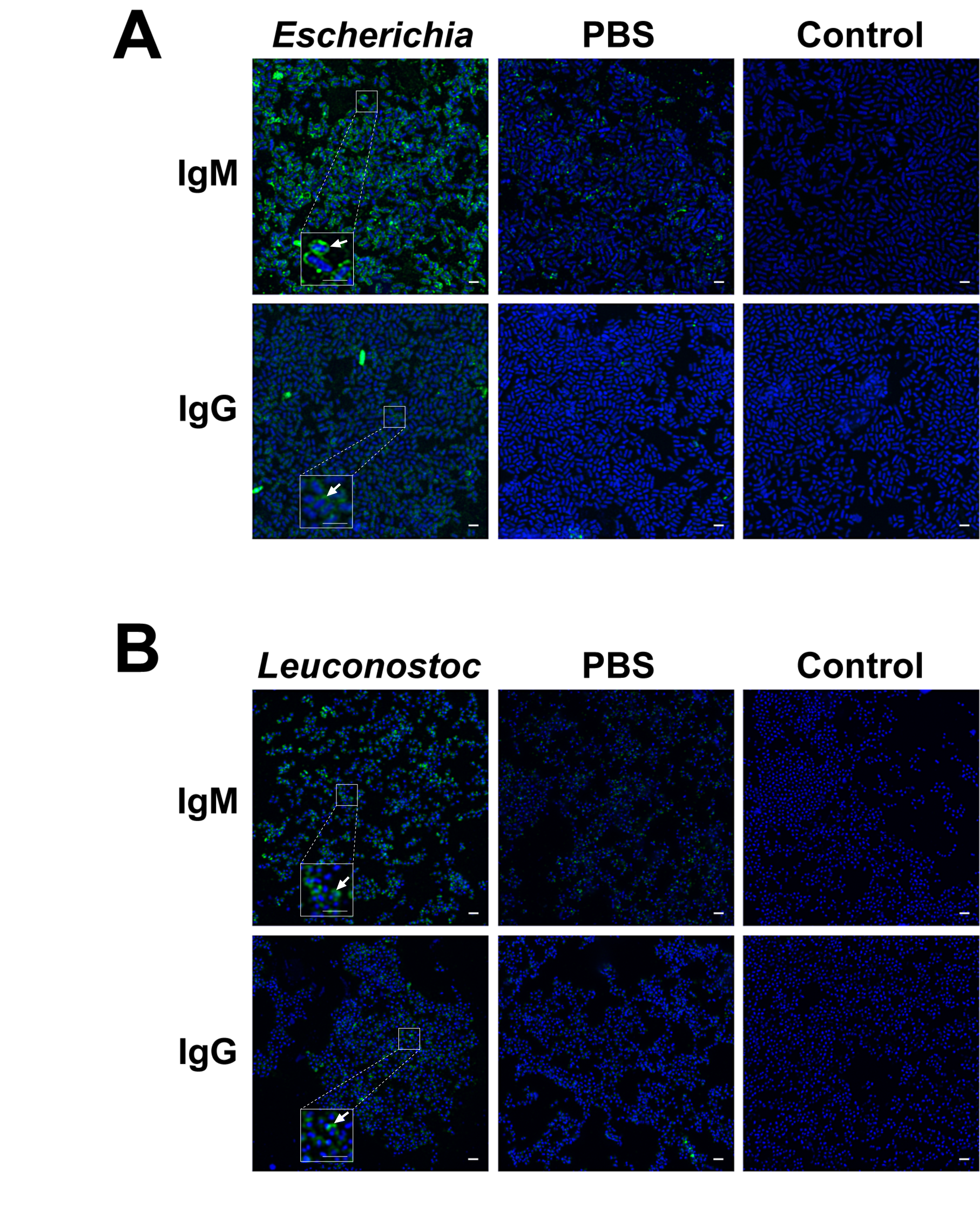

### Supplementary Figure S2

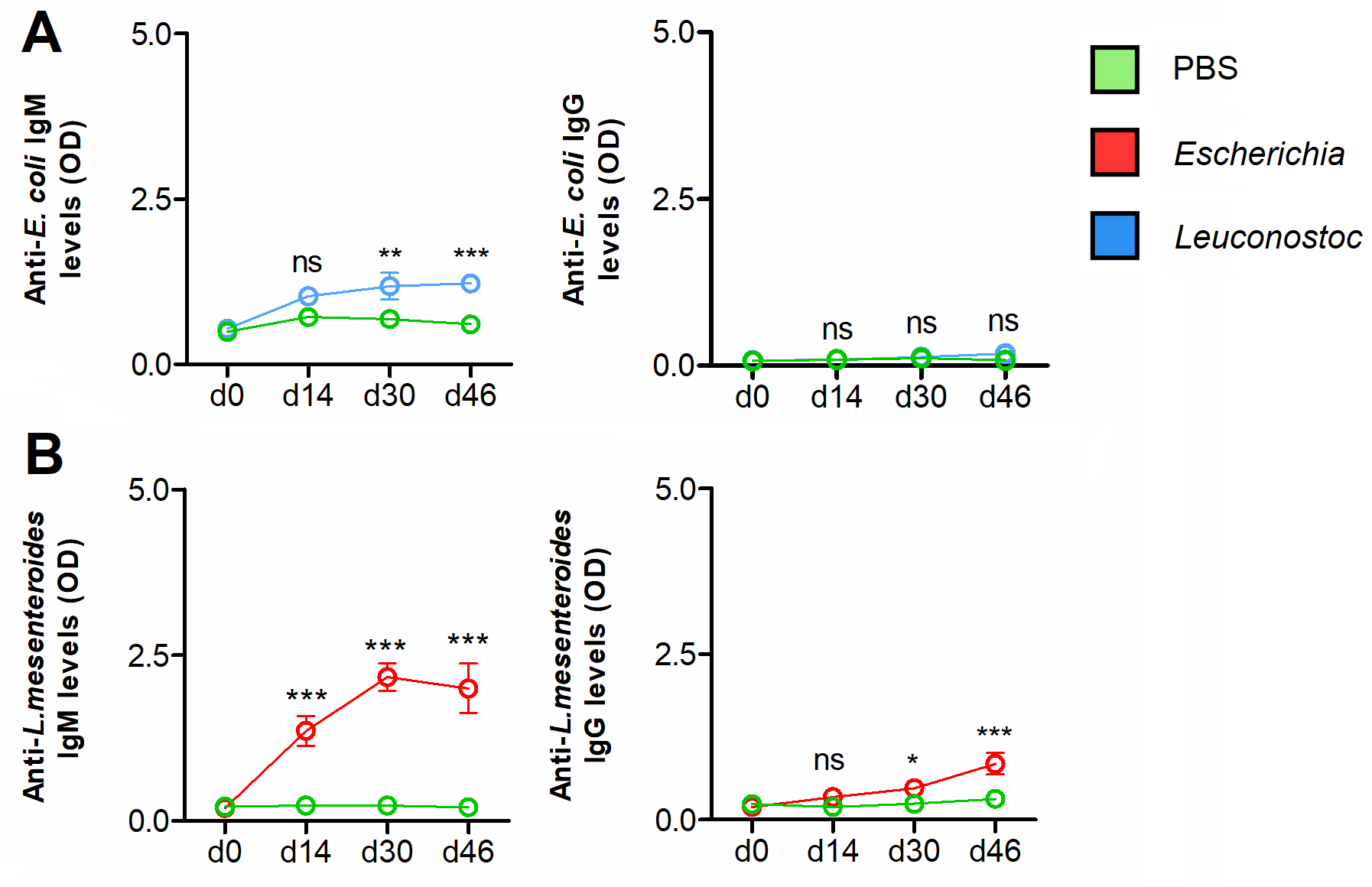

### Supplementary Figure S3

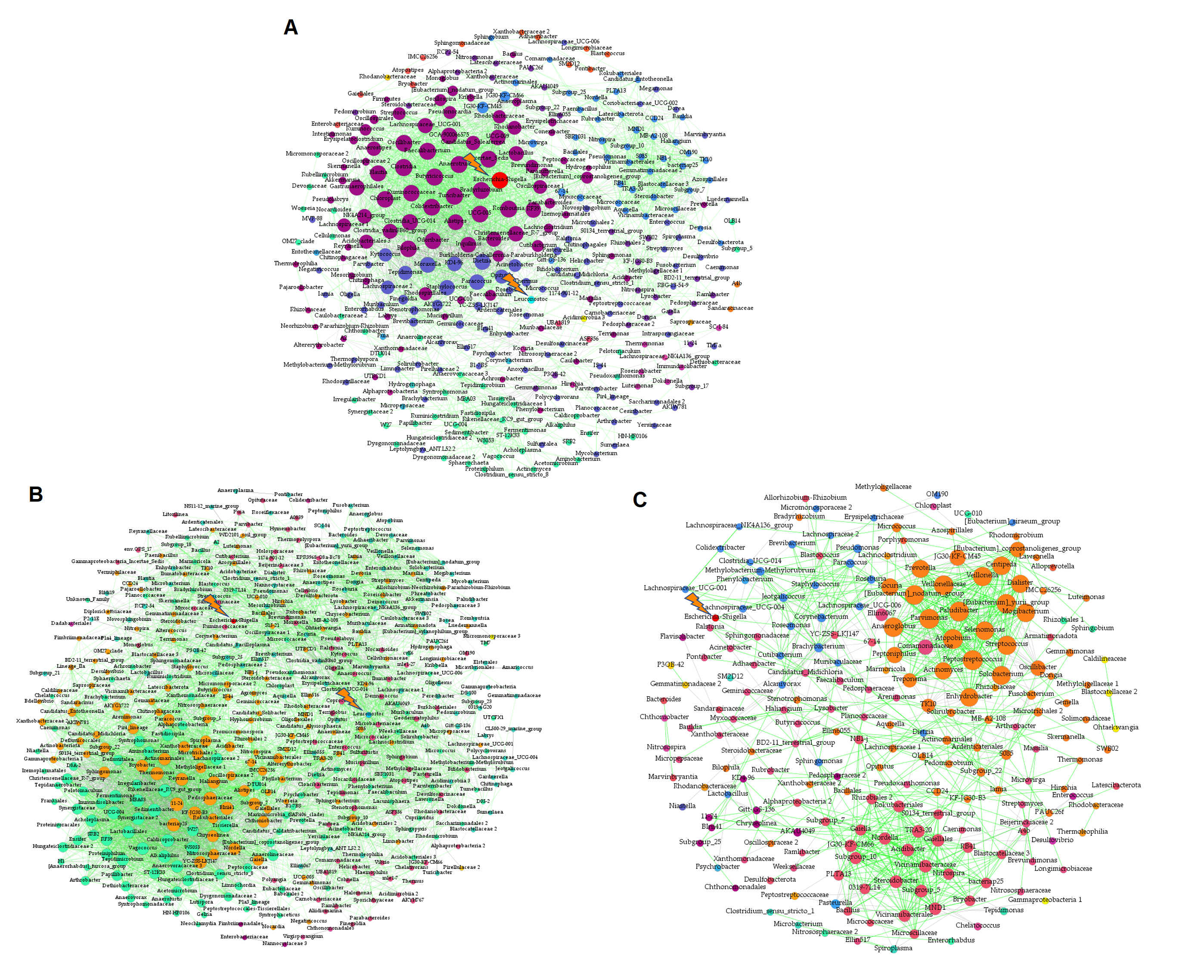
